# A new method for distinguishing human and mouse cells *in situ*

**DOI:** 10.1101/2020.06.08.139204

**Authors:** Yongqiang Wang, Zixian Huang, Kaishun Hu, Jiangyun Peng, Weicheng Yao, Weixi Deng, Jiyuan Zuo, Yin Zhang, Dong Yin

## Abstract

The mouse xenograft model is one of the most widely used animal model for biomedicine research. It is vital to distinguish the cells from different species, especially for the spatial distribution information. However, the available strategies of species-specific detection are either inapplicable *in situ* or of low specificity. Here, we reported a method based on DAPI staining, which offers an effective, convenient way that accurately identifies human and mouse nuclei at single-cell level *in situ*. This method was proven to be effective in cell co-culture and tumor xenograft tissue section. Microscopic imaging results shows obvious DAPI plaques-like structures in mouse nuclei, but absent in human nuclei. Moreover, we found these structures are co-localized with mouse major satellite DNA, which is located pericentromere in mouse, but absent in human. Our study provides a high-performance method that can be widely used for distinguish human and mouse cell *in situ*.

## Introduction

Animal models play critical roles in the field of biomedicine research, and the most commonly used is a mouse-based experimental animal model. Since the emergence of immunodeficient mice, mouse models for constructing human-mouse fits by transplanting cells or human tissues have been used widely, including cell line-derived xenografts (CDX) and patient-derived xenografts (PDX) Model(Olson, Li et al., 2018). It is of great significance to tracking transplanted cells and understanding the spatial relationship between donor and host cells. To distinguish cell sources in mouse xenograft model at the morphological level, antibody recognition methods (immunohistochemistry and immunofluorescence), transgenic reporter methods, or chromosome-specific probe labeling methods are widely used(Siracusa, Chapman et al., 1983; Démarchez M et al., 1993 ; Jacobsen PF et al., 1994; Matsuo S et al., 2007; Waldmann J et al., 2018). However, these methods all need to consider the issues of label specificity, signal intensity, and complexity of experimental operations, which increase the difficulty in use. This article reports a simple method for accurately distinguishing human and mouse nuclei based on DAPI staining. This method can not only accurately identify at the level of cultured cells, but also at cell properties in mouse xenograft model. Based on the short-wavelength excitation and high brightness of DAPI, and the simplicity of experimental operation, this method is expected to be widely used in the field of biomedical research.

## Results and discussion

### Human and mouse nuclei show significant morphological differences in response toDAPI staining

DAPI is a DNA-specific marker and is widely used in the field of life sciences (Kapuscinski,1995). It was interesting to investigate whether there was different pattern of DAPI-stained nuclei between human-derived cells and mouse-derived cells via a variety of fluorescence tests. Specifically, the cells displayed significant morphological differences. The nucleus of mouse-derived cells showed more obvious DAPI plaques, but this observation was absent in human-derived nucleus (Figure 1A and Supplementary Table 1). We counted the number of DAPI plaques in the nucleus of different tumor cells, including breast cancer, oral cancer, liver cancer, and primary macrophages derived from human and mouse (Figure 1B), and found that significant differences of DAPI plaques were existed in those cells. To confirm the differences of DAPI-staining of human and mouse nuclei in tissue samples, we further conducted detection in human and mouse breast, brain, lymphoid, liver, and intestine tissues. As expected, the similar results were observed in indicated tissues (Figure 1C). Those data insisted that there are distinct DAPI plaques in mouse cells but not in human cells, and this feature is irrelevant of cell lineages. In order to further confirm the specificity of DAPI staining to distinguish human and mouse cells at the single cell level, we constructed stable strains of mCherry-labeled or GFP-labeled human and mouse cells, respectively. As expected, two types of cells could be easily distinguished (4T1 cells and Hela cells)upon co-culture of these cells(Figure 2A, B, D) according to the feature of DAPI fluorescence plaques.

**Figure 1.**
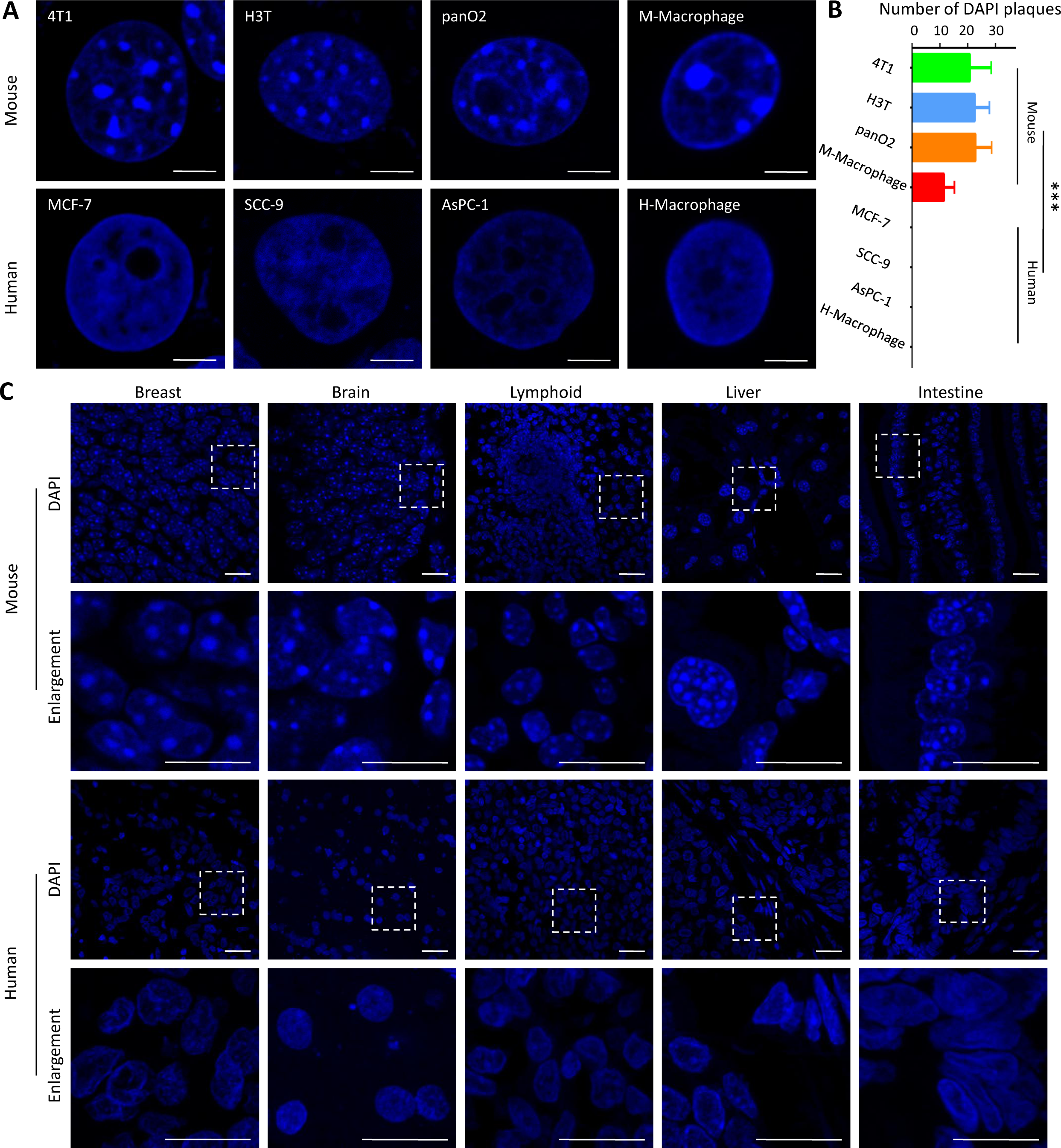
There were significant morphological differences between human and mouse nuclei upon DAPI staining. (A) DAPI staining was performed on four different types of cells derived from human and mouse. Scale bars, 5 μm. (B) DAPI plaques count analysis was performed on the cells indicated in (A). Asterisks (*) indicate a significant difference from the Human cells by *t*-test: *** *P* <0.01. (C) DAPI-labeled observations were performed on five different tissues derived from humans and mice, respectively. Scale bars, 20 μm.

**Figure 2.**
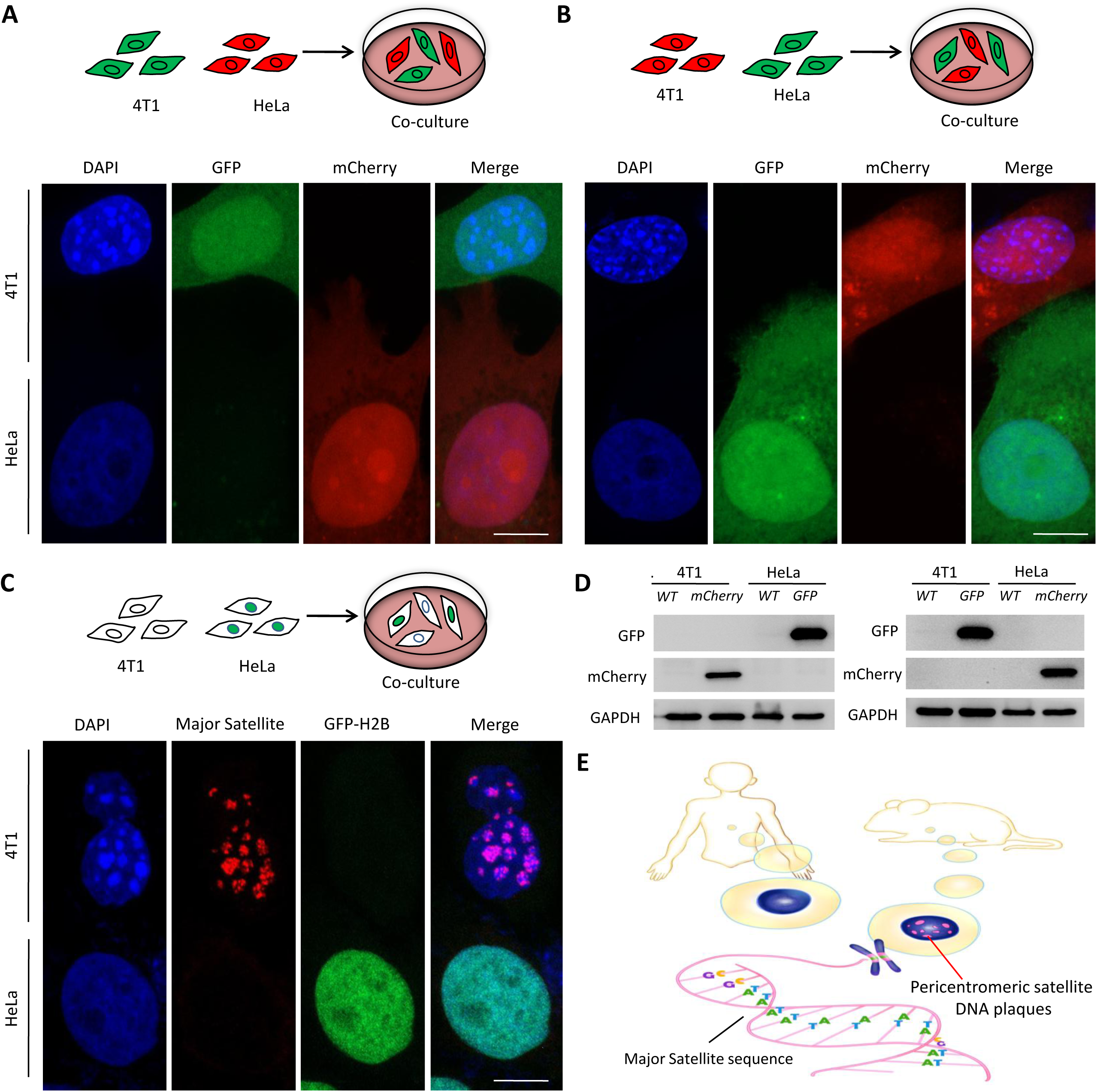
Identification of underlying mechanisms of DAPI plaques difference between human (HeLa) and mouse (4T1) cells.(A, B) The DAPI fluorescence feature can be clearly distinguished between human and mouse cells in the co-culture system. (C) Specific DAPI plaques in mouse nuclei co-localized with mouse major satellite fluorescence. (D) Immunoblot verification of fluorescent proteins indicated in (A, B) experiments. (E) Analytical model for morphological differences between human and mouse nuclear DAPI markers. Scale bars, 10 μm.

### The satellite DNA sequence in the mouse nucleus is the major determinants for distinguish mouse cells from human-derived cells upon DAPI staining

Previous studies have found that DAPI was a DNA-specific dye, which is mainly embed in (A + T)-rich regions(Zeman & Lusena, 1975). The distinct DAPI plaques in the mouse nucleus suggested that there were high-density heterochromatin-like regions in the DNA sequence of mouse nucleus, which were enrich in (A + T) sequences. The pericentromeric heterochromatin region of mice contains numerous non-coding satellite DNA sequences, which are rich in A-T sequences (Lyon MF& Searle AG, 1989;Jagannathan, Cummings et al., 2018). Therefore, we speculated that the main satellite DNA sequence of mouse centromeres might be the main reason for larger plaques in DAPI-staining mouse nuclei.

To verify the assumption, we employed a fluorescent probe target to mouse major satellite DNA sequence and verified it by fluorescence in situ hybridization (FISH). Mouse cells (4T1 wild-type) and human cells (GFP-H2B -labeled HeLa cell) were co-cultured for 24 hours. Then, a DNA hybridization experiment was performed. As showed in Figure 2C,the major satellite DNA sequence co-localized with DAPI plaques in mouse (4T1) cells, while there is no obvious FISH signal in human (Hela) cells.

These data suggested that DAPI can distinguish between human and mouse-derived cells due to the large number of major satellite DNA sequences in the nucleus of mouse-derived cells. These sequences generate specific plaques appear upon DAPI staining in mouse-derived cells. The human nucleus lacked this sequence, so no plaques appeared (Figure 2E). This observation makes it possible to distinguish between human and mouse-derived cells.

### Morphological differences in nuclei upon DAPI staining can accurately discriminate human and mouse cells in human-mouse xenograft model

Mouse models, especially CDX or PDX models, were one of the indispensable experimental methods in biomedical research. For the study of the cancer cells breaking through of tissue boundaries, the finding in this article has practical significance, which can clearly distinguish human tumor cells from normal mouse cells. Mouse subcutaneous tumor formation and tumor xenograft in situ models were applied in the following studies as the examined objects. From the experimental results in this study, we observed that there are significant DAPI plaques in the nucleus of mouse tissue cells, but no obvious plaques in the nucleus of human breast cancer tissues (Figure 1C and Figure 3A, B). We next performed method validation in a mouse PDX model. We found that there were two types of cells with significantly different DAPI morphology in tumor sections of PDX mice. One group of the nucleus had a significant DAPI plaques and the other group did not exhibit this phenomenon (Figure 3C). According to our finding, the cell with DAPI plaques belong to mouse nuclei, while the other nuclei belong to human-derived cells. We have confirmed this conclusion using in situ hybridization experiments with mouse major satellite sequences (Figure 3). Consistently, we have positively confirmed our method in mouse breast cancer CDX model (Supplementary Figure 1) and mouse liver cancer CDX model (Supplementary Figure 2).All the relevant tissues of the mouse models mentioned above were verified by H&E staining (Supplementary Figure 3).

**Figure 3.**
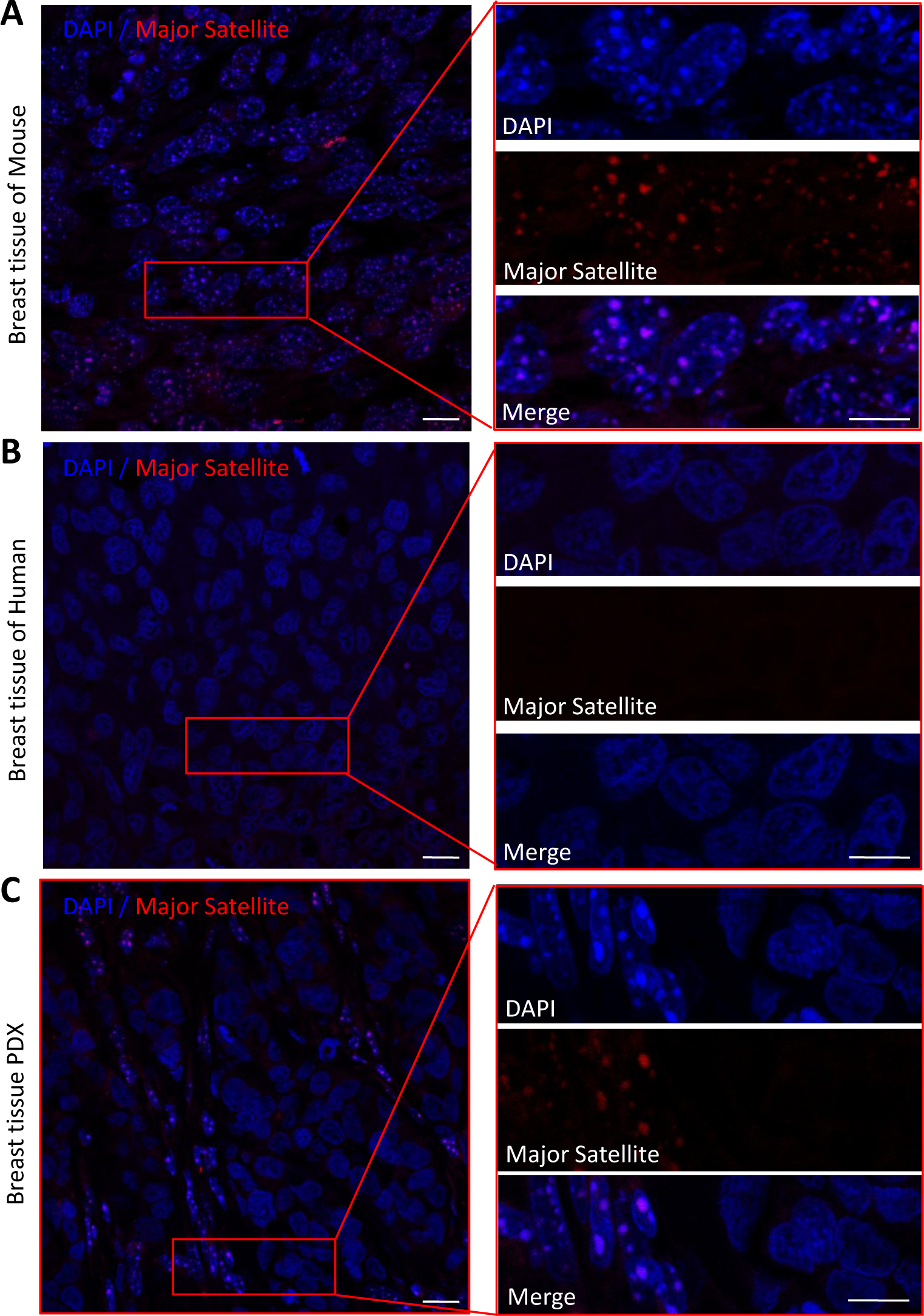
Validation of the reliability of human and mouse cell differentiation methods in breast cancer PDX models. (A) DAPI labeling in mouse breast cancer tissues and fluorescence verification in mouse major satellite. (B) Human-derived breast cancer tissue-level DAPI labeling and mouse major satellite fluorescence-negative verification. (C) DAPI labeling in mouse PDX model and fluorescence verification of mouse major satellite. Scale bars, 10 μm.

In this work, we found that the structure of human and mouse nuclei is significantly different in response to DAPI staining. The difference can accurately distinguish human and mouse nuclei at the level of co-culture cells and tumor xenograft tissue. Further experiments proved that the difference in nuclear DAPI coloration was related to the mouse genome centromere satellite DNA sequence. Based on this finding, we speculated that any DNA-specific marker can show the difference between the two types of nuclei, including Hoechst #33258(Moser, Dorman et al., 1975; Lawrence et al., 1977; Cunha & Vanderslice, 1984; Kozak, Miller & Ferrara, 1988), Hoechst #33342 (data not shown) and histone (GFP-H2B) Specific fluorescent labeling (Kanda, Sullivan et al., 1998); Supplementary Figure 4), etc. In the future, the aspects related to human and mouse cells can be considered for further application through this method, such as drug development, cell identification, identification of true and false fluorescent pictures, etc.

## Methods

### Cell culture

Both mouse and human cell lines used in this work were obtained from the American Type Culture Collection (ATCC), and cultured in DMEM or F12 medium supplemented with 10% FBS. Macrophages were obtained by referring to literature and cultured in complete medium containing serum(Baghdadi, Wada et al., 2016; Weischenfeldt & Porse, 2008). All the cells were cultured at 37 °C humidified incubator containing 5% CO_2_.

### Mouse model and tissue acquisition

Local tumor tissues removed from breast tumor patients were used to establish a mouse model of PDX. The patient’s written informed consent was obtained in advance, and the research protocol was approved by the hospital ethics committee. All experiments used immunodeficient mouse were performed according to guidelines approved by the Institutional Animal Health and Use Committee (IACUC). Fresh surgical tumor tissues were implanted directly into immunodeficient mouse to establish a PDX mouse model, and SNP48 analysis confirmed that the established PDX was successful. In the breast tumor cell line transplant model, we used MCF-7 cells. The cultured cells were implanted directly into the skin of immunodeficient mouse to establish a CDX mouse model. In the orthotopic transplantation model of liver tumor cell lines, we used Huh7 cells, and the cultured cells were directly implanted into the liver of immunodeficient mouse in situ to establish a mouse model. After tumor masses were formed in the above mice, they were extracted and embedded in paraffin for sectioning.

### DNA fluorescence in situ hybridization

The prepared cell and tissue samples were fixed with 4% formaldehyde for 10 minutes, and then washed in PBS-T for 30 minutes. The fixed sample was incubated with 2 mg / ml RNase A solution at 37 ° C for 10 minutes, and then washed with PBS-T +1 mMEDTA. The samples were then washed in 2xSSC-T (2xSSC containing 0.1% Tween-20) for 15 minutes with increasing formamide concentration (20%, 40% and 50%), and finally washed in 50% formamide for 30 minutes. Probe hybridization buffer (50% formamide, 10% dextran sulfate, 2xSSC, 1 mMEDTA, 1 mM probe) was added dropwise to the washed samples. The samples were denatured at 91 °C for 2 minutes and then incubated at 37 °C overnight. After washing 3 times with 2xSSC, staining with 10 μg / ml DAPI for 10 minutes, and finally washing with PBS three times and mounting samples for observation. Probe used: major satellite-5’-Cy3-GGAAAATTTAGAAATGTCCACTG-3’(Jagannathan et al., 2018).

### Fluorescence microscopy

Confocal micrographs were acquired on a Zeiss LSM800 Laser Scanning Microscope equipped with a Plan Apochromat × 40 / 1.4 NA oil immersion objective. The photo shooting used the Z-STACK mode of ZEN software uniformly, and then uses the orthogonal projection to display multiple layers of signals after superposition. Areas with significant DAPI fluorescence intensity in the nucleus (about 2-3 times the fluorescence intensity in other areas of the nucleus) were recorded as plaques, and the number of plaques in 20 nuclei was calculated for each kind cell for statistics

### Western Blot

For protein extraction, the cells were washed twice with cool phosphate-buffered saline (PBS), harvested by scraping and then lysed in lysis buffer (#9803, CST). Following centrifugation, the supernatant was collected, and the protein concentration was determined using the BCA Protein Assay Kit (#23227, Thermo Scientific).

For Western blotting, cell lysates were electrophoretically separated on an SDS-PAGE gel using a standard protocol. The proteins were then transferred to Immobilon-P transfer membranes (PVDFs) (IPVH00010; Millipore). The membranes were blocked with 5% non-fat milk in Tris-buffered saline containing 0.1% Tween-20 (TBST) for 1 hour at room temperature. The blots were incubated with the antibodies (mCherry #43590, CST and GFP #2956, CST) at 4°C overnight, washed in TBST and then probed with the appropriate secondary antibody. Western blot analysis was performed according to standard protocols.

## Acknowledgments

This work was supported by grants from the Natural Science Foundation of China [81872140, 81420108026, 81572484, 81621004 to D.Y., 31801075 to Y.Z.]; Guangdong Science and Technology Department [2019B020226003 to D.Y., 2018A030310344 to Y.Z. 2017B030314026]; Guangzhou Bureau of Science and Information Technology [201704030036 to D.Y.]; Fundamental Research Funds for the Central Universities [18zxxt62 to Y.Z]

## Author contributions

Yongqiang Wang, Zixian Huang, Kaishun Hu, Data curation, Formal analysis, Methodology; Jiangyun Peng, Weicheng Yao, Weixi Deng, Data curation, Formal analysis; Jiyuan Zuo, Data curation; Yin Zhang, Supervision, Funding acquisition, Writing - review and editing; Dong Yin, Conceptualization, Supervision, Funding acquisition, Writing - review and editing.

## Competing interests

The authors declare that they have no competing interests.

## Notes

### Competing Interest Statement

The authors have declared no competing interest.

