## Supplemental Figure and Table for "A new method for distinguishing human and mouse cells *in situ*"

### Supplementary Material

**Supplementary Table 1.** Some human and mouse cells verified in this work.

**Supplementary Figure 1.** Validation of the reliability of human and mouse cell identification methods in a breast cancer CDX model (the cell line of this experiment was MCF-7). Scale bars, 10 μm.

**Supplementary Figure 2.** Validation of the reliability of human and mouse cell differentiation methods in a liver cancer CDX model. (A) DAPI labeling at mouse liver cancer tissue level and fluorescence verification of mouse major satellite. (B) Human-derived liver cancer tissue level DAPI labeling and mouse major satellite fluorescence negative verification. (C) DAPI labeling in mouse CDX model and fluorescence verification of mouse major satellite (the cell line of this experiment is Huh7). Scale bars, 10 μm.

**Supplementary Figure 3.** H&E staining was conducted to verify mouse xenograft tissue sections. (A) HE staining of human and mouse tissue sections related to breast cancer. (B) HE staining of liver cancer-related human and mouse tissue sections. Scale bars, 25μm.

**Supplementary Figure 4.** Histone H2B fluorescence (green) and DAPI have similar marking effects on nuclear chromosomes. Scale bars, 10 μm.

### Supplementary Table 1

| | Mouse | Human |
|---|---|---|
| Cell | 4T1, H3T, panO2, Mouse Primary Macrophages, hepa1-6, Natural killer cell, RAW264.7, Mouse Primary Fibroblasts, etc. | MCF-7, SCC9, AsPC-1 , Human Primary Macrophages, UM-UC-3, t24, HepG2, MDA-MB-231, MHCC97-H, PC-3, BT474, HCCLM3, HuH7, THP-1, SiHa cell, HuH-6, etc. |

### Supplementary Figure 1

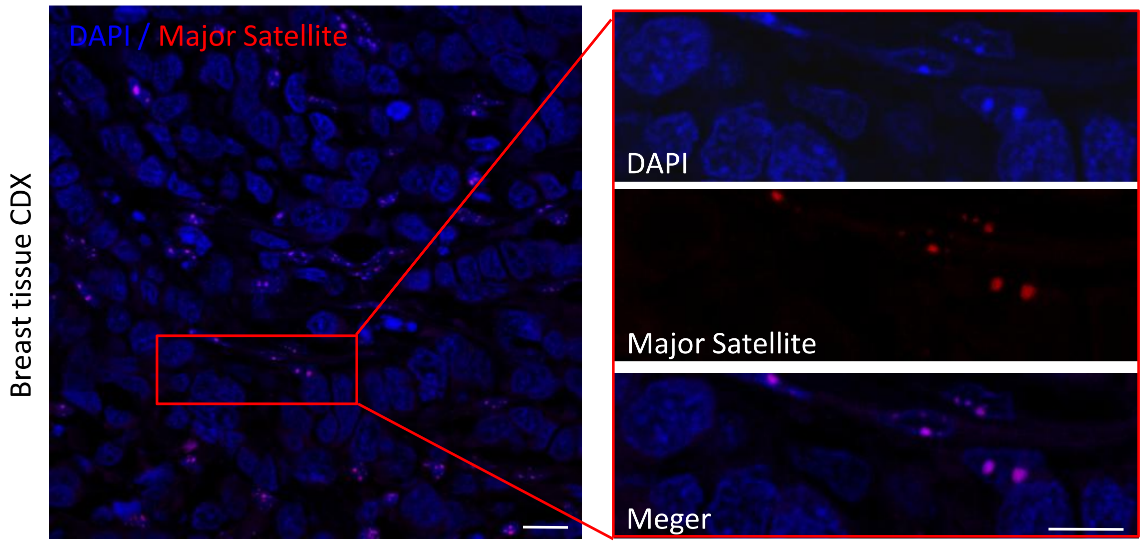

### Supplementary Figure 2

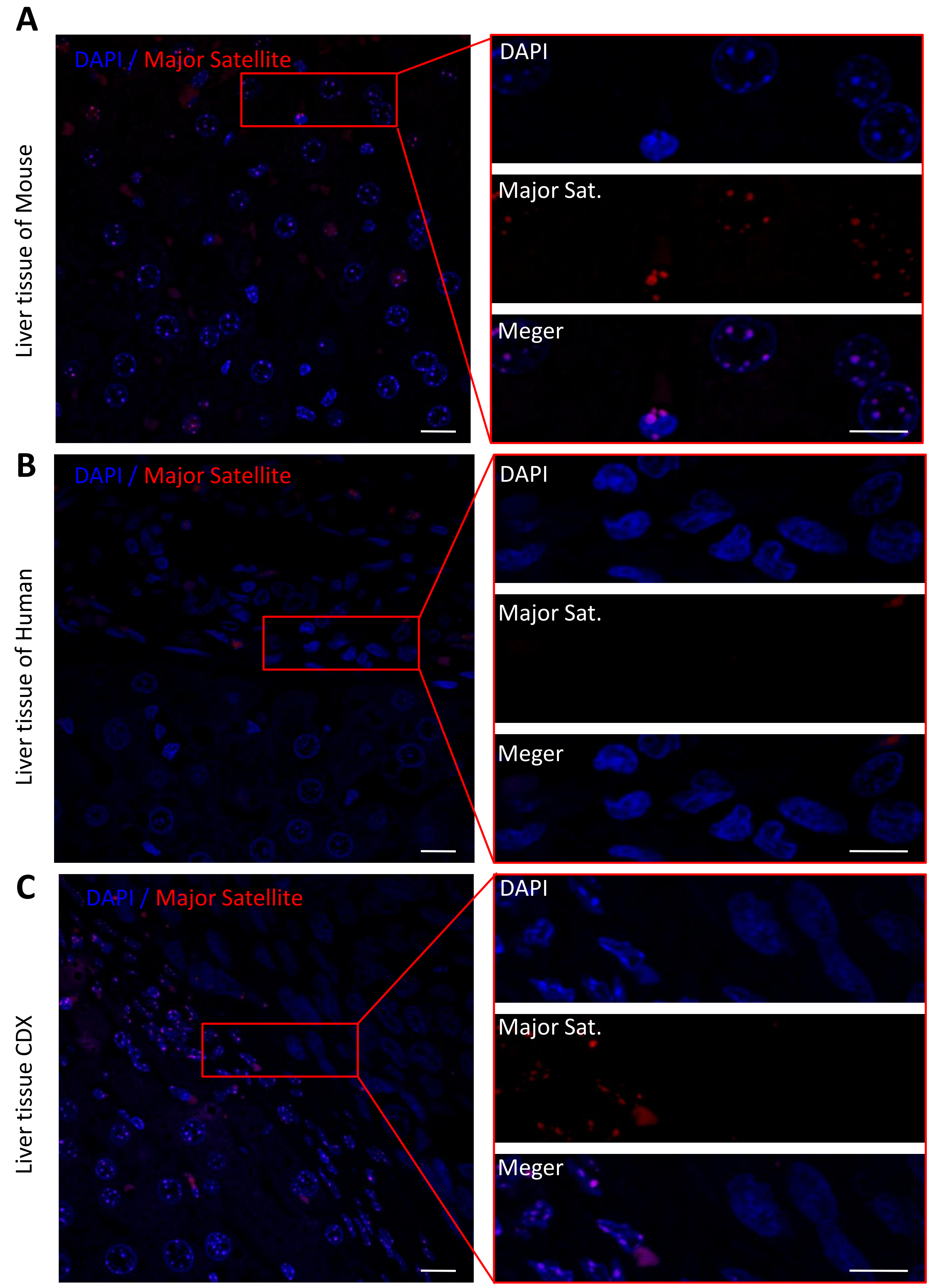

### Supplementary Figure 3

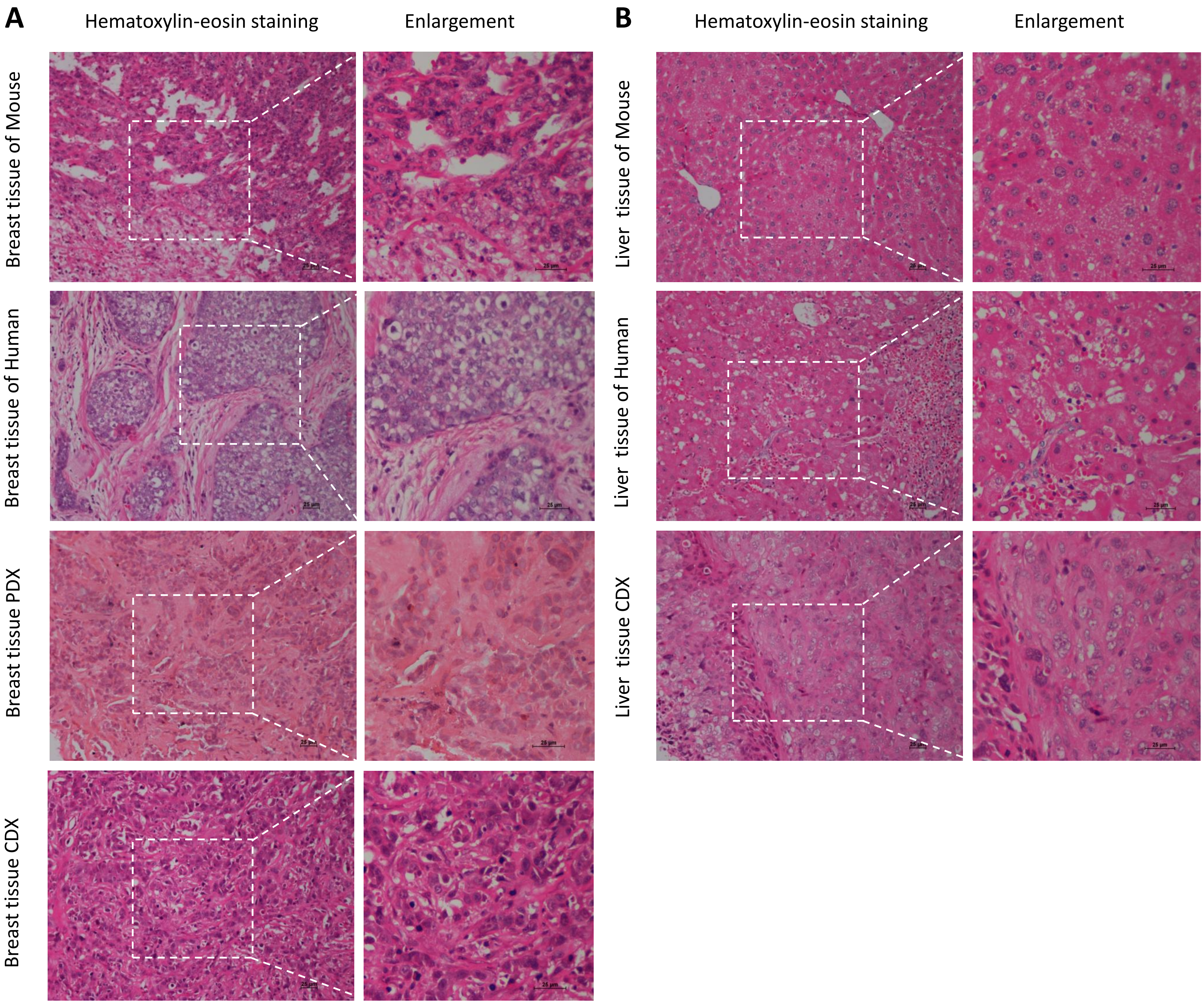

Supplementary Figure 4

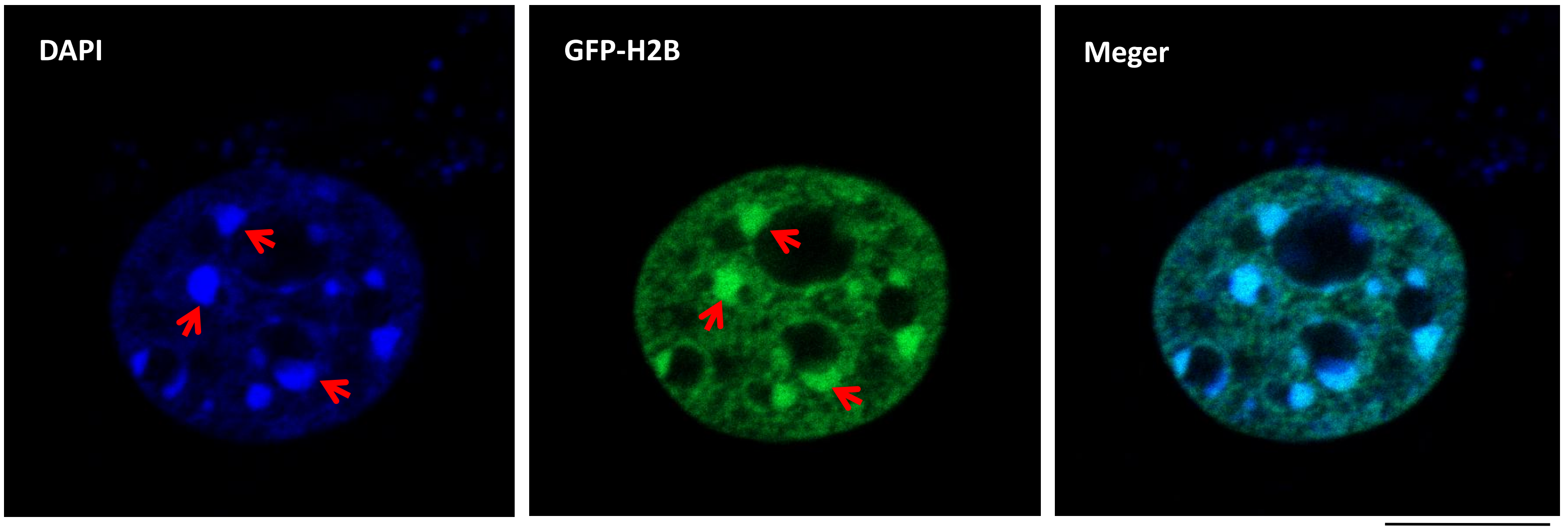
